# Systematic identification of transcriptional activator domains from non-transcription factor proteins in plants and yeast

**DOI:** 10.1101/2023.09.12.557247

**Authors:** Niklas F. C. Hummel, Kasey Markel, Jordan Stefani, Max V. Staller, Patrick M. Shih

## Abstract

Transcription factors promote gene expression via trans-regulatory activation domains. Although whole genome scale screens in model organisms (e.g. human, yeast, fly) have helped identify activation domains from transcription factors, such screens have been less extensively used to explore the occurrence of activation domains in non-transcription factor proteins, such as transcriptional coactivators, chromatin regulators and some cytosolic proteins, leaving a blind spot on what role activation domains in these proteins could play in regulating transcription. We utilized the activation domain predictor PADDLE to mine the entire proteomes of two model eukaryotes, *Arabidopsis thaliana* and *Saccharomyces cerevisiae* (*1*). We characterized 18,000 fragments covering predicted activation domains from >800 non-transcription factor genes in both species, and experimentally validated that 89% of proteins contained fragments capable of activating transcription in yeast. Peptides with similar sequence composition show a broad range of activities, which is explained by the arrangement of key amino acids. We also annotated hundreds of nuclear proteins with activation domains as putative coactivators; many of which have never been ascribed any function in plants. Furthermore, our library contains >250 non-nuclear proteins containing peptides with activation domain function across both eukaryotic lineages, suggesting that there are unknown biological roles of these peptides beyond transcription. Finally, we identify and validate short, ‘universal’ eukaryotic activation domains that activate transcription in both yeast and plants with comparable or stronger performance to state-of-the-art activation domains. Overall, our dual host screen provides a blueprint on how to systematically discover novel genetic parts for synthetic biology that function across a wide diversity of eukaryotes.

**Significance Statement:** Activation domains promote transcription and play a critical role in regulating gene expression. Although the mapping of activation domains from transcription factors has been carried out in previous genome-wide screens, their occurrence in non-transcription factors has been less explored. We utilize an activation domain predictor to mine the entire proteomes of *Arabidopsis thaliana* and *Saccharomyces cerevisiae* for new activation domains on non-transcription factor proteins. We validate peptides derived from >750 non-transcription factor proteins capable of activating transcription, discovering many potentially new coactivators in plants. Importantly, we identify novel genetic parts that can function across both species, representing unique synthetic biology tools.

## Introduction

Transcription factors (TFs) regulate gene expression by binding specific DNA regions with their DNA-binding domains (DBDs) and interacting with protein complexes through their transcriptional effector domains (2). Transcriptional effector domains that promote transcription are further classified as activation domains (ADs). In this work, we will refer to all short protein sequences that function in AD assays as ADs, regardless of their native function. New high-throughput methodologies have helped characterize the regulatory activity of transcriptional effector domains *en masse* in yeast, human, and fly models (3–9), and these approaches are beginning to be implemented to study plants (10). Still, most studies have largely biased their focus on TFs, leaving other nuclear proteins and their potential role in transcription understudied. Moreover, non-nuclear proteins have been demonstrated to be involved in transcriptional regulation. For example, Notch1, a plasma membrane localized protein in multicellular animals, contains a C-terminal AD which is cleaved, processed, and localized to the nucleus to induce transcription (11). Similarly, the cell-adhesion protein beta-catenin is localized to the nucleus when multimerized where it acts as a transcriptional coactivator in fly and vertebrates and the closest plant homologs have been linked to root development (12–14). Thus, there is evidence of proteins with ADs outside of standard TFs. To more thoroughly study all putative proteins that may be involved in transcriptional activation, genome wide screens of all proteins – not just TFs, or nuclear proteins – are needed to identify previously unannotated molecular factors that may play a role in transcriptional regulation.

The availability of large AD activity datasets has enabled the development of deep convolutional neural networks that can predict the activity of eukaryotic ADs from protein sequences (1, 6). These models have helped elucidate how specific amino acid sequence features of acidic ADs enable their transcriptional activation activity (15). Importantly, because current AD predictive models have been trained on large datasets from select organisms (i.e., yeast and human), the predictive strength of these models in other eukaryotes has not been well defined. The published neural network models have never been directly tested in a large-scale experiment.

Mechanistic studies have shown how acidic residues promote the exposure of hydrophobic residues that in turn are essential for AD activity (16). Hence, the distribution of acidic and hydrophobic residues is key, as hydrophobic clusters can lead to the intramolecular collapse of the AD, diminishing its activity (7). The recently proposed acidic exposure model links these observations to structural disorder in ADs, where acidic residues stabilize an energetically unfavorable solvent exposure of hydrophobic residues which in turn interact with coactivators to promote transcription in a transiently structured fashion (7). Thus, sequence composition, structural disorder and small sequence motifs in ADs have been linked to defining AD activity, but we still lack a comprehensive understanding of how positional sequence features affect AD function.

Eukaryotic transcription is facilitated by TFs, coactivators, and chromatin regulators. Coactivators can function as adaptors between TFs and RNA polymerase II or the general transcription apparatus, while other coactivators modify chromatin to help transcription of chromatinized templates or help with unwinding DNA, all resulting in higher transcriptional output (11–14, 17–19). Coactivators interface between TFs and RNA polymerase but do not directly bind DNA, functionally separating them from TFs. Coactivators and chromatin regulators can contain ADs (1, 8, 20), marking activator activity non-unique to TFs; still, there has been a dearth of high-throughput studies focused on identifying new coactivator candidates, due to the multitude of mechanisms that coactivators use to promote transcription. Hence, the occurrence of ADs in nuclear non-TF genes could indicate that a given protein is involved in transcription and help annotate previously unknown transcription associated genes and coactivators.

Genome-scale studies characterizing TFs in plants have provided the foundational understanding of the complex regulation that underlies plant development, adaptation, and overall physiology (21, 22). However, transcriptional coactivators have been much less studied and leaves a large blind spot in our understanding of their role in transcription. Unlike unicellular systems that are more readily tractable to screening massive libraries and cell sorting, the complex physiology and cell wall of plants has hindered the implementation of high-throughput methods for the characterization of ADs in plants. As a result, our understanding of plant ADs and the role of potential coactivators pales in comparison to other better-studied model eukaryotes (e.g. yeast, human, etc.). We previously reported that a machine learning model trained on data from a large library of synthetic activators from yeast can correctly predict ADs in plant TFs (10); however, it is still unclear how applicable and scalable these models are in plant systems, necessitating further evaluation of more plant ADs predicted by yeast models. A larger set of validated plant ADs would allow the comparison of sequence features in plant ADs with observations in other well-studied eukaryotes. Moreover, studying ADs from non-TF genes can help us identify and annotate previously unknown proteins involved in transcriptional regulation and deepen our understanding of the features defining AD strength in plants.

Here, we assess the transcriptional activity of predicted ADs derived from non-TF proteins from yeast and plants. We generated a library of 18,000 synthetic TFs carrying predicted ADs from non-TF genes with ADs derived from *S. cerevisiae* and *A. thaliana*. We show that 753 (89%) of 846 parent genes in the library contain ADs capable of promoting transcription in yeast, identifying many proteins previously unassociated with the activation of transcription. Notably, ADs were not limited to nuclear genes and many ADs were found in a wide range of protein families localized to other organelles. We find positional distribution of key amino acids that make large contributions to AD activity providing insight into sequence grammar. Furthermore, we show how strong ADs from the library activate transcription in plants, marking them as universal ADs. Our large interspecies dataset provides both the foundational knowledge to explore the role of ADs in non-TFs as well as a large set of new ADs that can be readily integrated into genome engineering efforts across phylogenetically diverse eukaryotes.

## Results

### Characterization of a library of non-TF ADs mined from yeast and plant proteomes

We aimed to systematically discover previously uncharacterized ADs derived from non-TF proteins in two model eukaryotic systems, *A. thaliana* and *S. cerevisiae*. In previous work, we have shown that AD predictors derived from fungal data can accurately predict ADs in plant TFs and that plant ADs function in yeast (10). Here, we leveraged this result to predict activators in plant and yeast proteins, followed by high-throughput experimental validation in yeast.

To extract potential ADs from both proteomes, we utilized PADDLE, a neural network model capable of predicting acidic ADs in 53 amino acid long peptides (1). We computationally chopped each proteome in 53 AA tiles spaced every one residue, yielding 9,211,910 tiles in *A. thaliana* and 2,646,422 tiles in *S. cerevisiae* derived from 27,082 and 6,455 proteins, respectively **(Fig. 1A, B)**. We used PADDLE to predict the potential of all tiles to activate transcription. We then used TF databases for both species – PlantTFDB v5.0 and Yeastract+ – to remove all tiles derived from TF sequences (23, 24). We found that tiles from non-TF genes had a similar dynamic range of predicted activity as tiles from TF genes **(Fig. S1A, B)** and in Arabidopsis the strongest predicted tile occurred in a non-TF gene (AT5G07570.1). We defined all genes that we mined tiles from as parent genes, as multiple tiles can come from a single protein sequence. We then selected 12,000 tiles from *A. thaliana* and 6,000 tiles from *S. cerevisiae* with the highest predicted activation score, yielding a 18,000 tile library derived from 447 Arabidopsis and 402 yeast parent proteins, respectively. We chose to include overlapping tiles to increase accuracy and resolution. To gauge the subcellular localization of parent proteins of the library, we utilized SUBA5 for Arabidopsis and YeastGFP/YPL+ to annotate localization (25–27). There were a total of 214 parent proteins localized to the nucleus in Arabidopsis and 107 in yeast. Non-nuclear genes were localized throughout all subcellular locations in both species **(Supplementary Table 1, 2)**. This diversity of localization suggests that peptides predicted to be ADs occur throughout various organelles across both proteomes.

**Figure 1.**
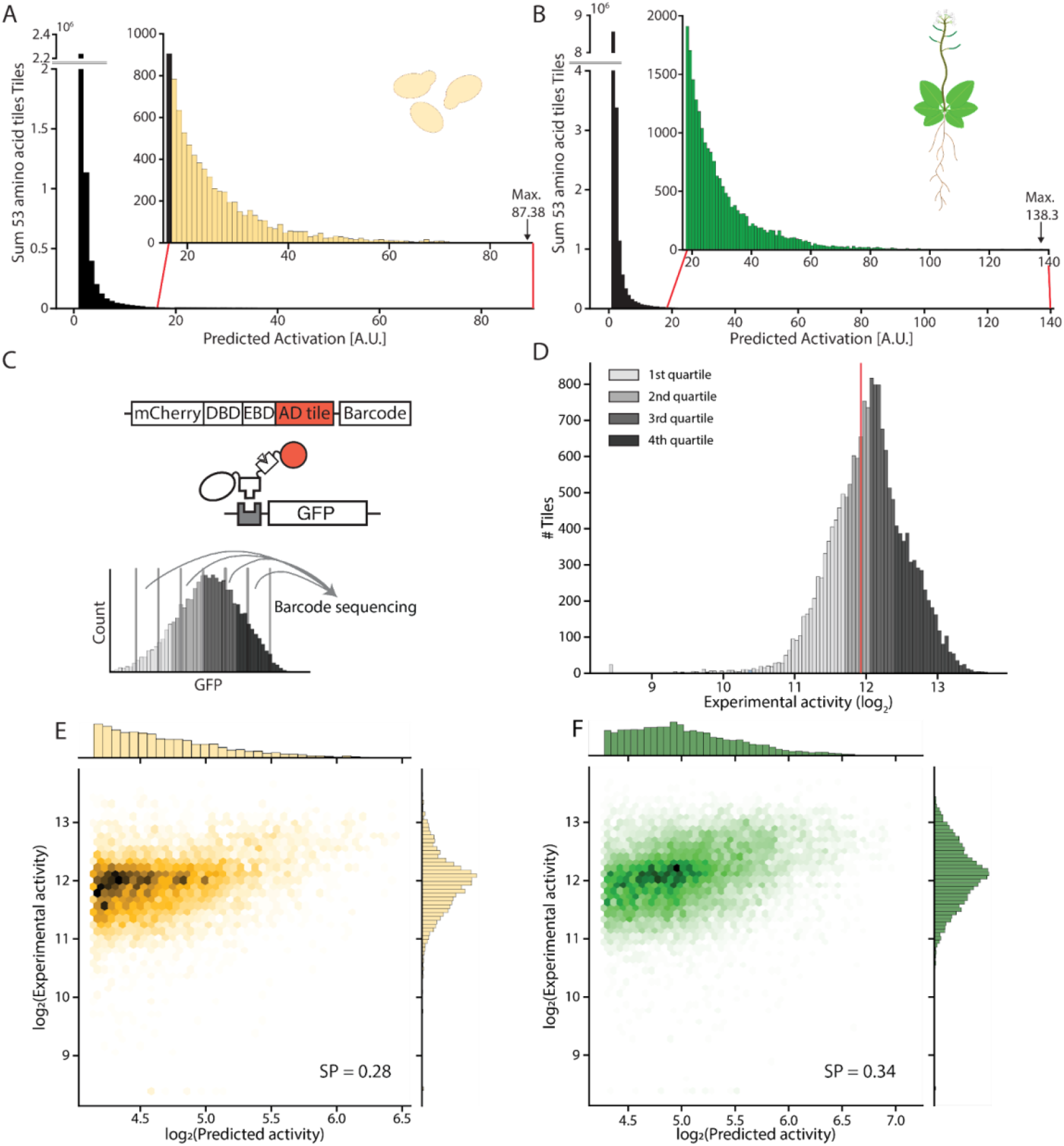
Proteome-wide characterization of putative ADs mined from non-TF plant and yeast proteins. Histogram showing the PADDLE predicted activity of all 53 amino acid tiles in the (**A**) *S. cerevisiae* and (**B**) *A. thaliana* proteome. Inlet figures show the magnified areas of the histogram the putative AD candidates for the libraries were chosen from (marked in red). (**C**) The 12,000 strongest *A. thaliana* and 6,000 strongest *S. cerevisiae* tiles were characterized as a synthetic TF library in *S. cerevisiae*. Activator activity was calculated by abundance of barcodes in bins established during FACS sorting. DBD: DNA binding domain, EBD: estrogen binding domain. (**D**) Activity of every tile as determined by FACS and consecutive barcode sequencing across eight bins. Red bar indicates activity of no-AD controls. Predicted activity vs. experimentally observed activity of all tiles from (**E**) *S. cerevisiae* and (**F**) *A. thaliana*. SP: Spearman’s R.

To experimentally characterize and validate our library, we used a previously established expression system utilizing synthetic TFs in yeast (16). In this expression system each tile is fused to a synthetic TF, consisting of 1) mCherry for normalization of TF concentration to generated reporter signal as a N-terminal fusion, 2) the orthogonal murine Zif268 DBD, 3) a human estrogen response domain to make the system inducible with ß-estradiol, 4) the 53 amino acid long AD candidate, and 5) a unique barcode in the 3’ UTR marking candidate identity in the library **(Fig. 1C)**. The associated reporter consists of six copies of the Zif268 binding sites upstream of a modified GAL1 promoter driving GFP (28). Both the reporter and the synthetic TF were integrated into the genome of *S. cerevisiae* to reduce expression variability. We used fluorescence-activated cell sorting (FACS) to sort the library (see Methods) and experimentally validated the activity of 17,553 tiles (97.5% of total library) from 846 parent genes with high reproducibility between replicates (Pearson’s r = 0.82) **(Fig. S2, Supplementary Table 3, 4)**. Multiple DNA barcodes were used for each tile to further measure the variability resulting from multiple integrations, thereby increasing accuracy of measurements as previously shown (16).

The experimental activity of our library allowed us to evaluate the accuracy of PADDLE predictions. In the PADDLE training dataset, ∼30% of TF derived tiles showed activity and the model achieved reliable qualitative and quantitative prediction of ADs (∼10400 tiles, Pearson’s r = 0.81). In our library, tile activity ranged over three orders of magnitude with 56.5% of the library showing significant activity above no-TF control levels **(Fig. 1D)**. This was the largest fraction of active tiles we have observed using this system (7, 16). Parts of the library activated transcription equally or stronger than Gcn4-AD and -VP16-AD controls **(Fig. S3A, B)**. We found 89.0% of parent genes (753 out of 846) of tiles in our library to contain at least one tile with activator activity, demonstrating that tiles that can function as ADs are widespread throughout non-TF genes. This result shows how PADDLE can in most cases correctly localize ADs in proteins, but that its architecture, which predicts AD likelihood and then extrapolates activity predictions, is not rigorous enough to predict both qualitative and quantitative aspects of ADs. Overall, we identified 9911 novel ADs derived from non-TFs, providing a rich resource for engineering efforts.

### Single amino acid tiling unravels positional effects and key residues dictating AD activity

While the large fraction of active tiles supports the ability of PADDLE to localize ADs in protein sequences, the quantitative predictions of AD strength did not correlate as strongly with our experimental results (Spearman’s r = 0.35, Pearson’s r = 0.33) **(Fig. S4)**. Notably, the PADDLE algorithm predicted activity with higher accuracy in *A. thaliana* than in *S. cerevisiae* AD populations with moderate Spearman correlation coefficient of 0.34 and 0.28, respectively **(Fig. 1E, F)**. We found that in 13% of parent genes the strongest predicted tile precisely overlapped with the strongest experimental tile. Hence, PADDLE correctly identifies the general location of ADs but struggles to accurately predict the quantitative strength of the respective AD, as well as precise AD boundaries. Our results support previous evidence that there are positional effects of amino acid residues dictating AD activity, and demonstrate that PADDLE cannot resolve these effects.

To further investigate the discrepancies between PADDLE predictions and observed activities, we examine how amino acid composition and positional context may play a role in defining AD activity. We recovered previously observed relationships between amino acid sequence and activity, i.e. that the W, F, L, D, and E are associated with activity, and K and R are not (1, 6, 29, 30). To compare tile populations, we split the entire library into four equal quartiles reaching from weakest to strongest activity. Notably, the average amino acid composition of tiles in all four quartiles was nearly identical **(Fig. S5)**. This result emphasizes the power of this dataset to probe sequence grammar. We found the enrichment of acidic-aromatic dipeptides in strong ADs was reproducible in our dataset **(Fig. S6A-D, Supplementary Data 1)** (**6**). We hypothesized that the positional distribution of functional amino acids is key to AD activity.

To gauge the positional information encoded in each quartile, we measured the local density of all residues along each tile of every quartile, where density is the frequency of the respective amino acid in a 5 amino acid window. We then grouped amino acid groups linked to AD activity and computed density at each position of the 53 amino acids of every tile of each respective quartile. We found the density of W, F, Y, L residues, which are closely linked to AD activity, to be overall higher in the highest activity quartile, and density was higher in the C-terminus when compared to tiles in other quartiles **(Fig. 2A, Fig. S7A)**. This finding supports the acidic exposure model, which predicts that hydrophobic residues at the C-terminus will be more exposed to solvent and make larger contributions to activity. All quartiles had a low density of hydrophobic residues in the N-terminus suggesting that PADDLE has partially learned this signal. Correspondingly, the fourth quartile displayed a weaker density of acidic residues in the C-terminus and was more evenly distributed throughout the entire tile, whereas the weaker quartiles had a stronger enrichment of acidic residues in the C-terminus and depletion in the N-terminus **(Fig. 2B, Fig. S7B)**. These results support the hypothesis that the occurrence of key amino acids – in this case hydrophobic and acidic residues – alone does not correlate with activity but rather their distribution along the AD, further supporting the acidic exposure model (7). The dip in acidic residues at the extreme C-terminus in the 4th quartile was surprising to us, because we and others have previously shown that acidic residues near or next to aromatic residues boost activity. We speculate that the C-terminus is highly exposed, both by virtue of being on the end and because of the additional acidity of the C-terminal backbone.

**Figure 2.**
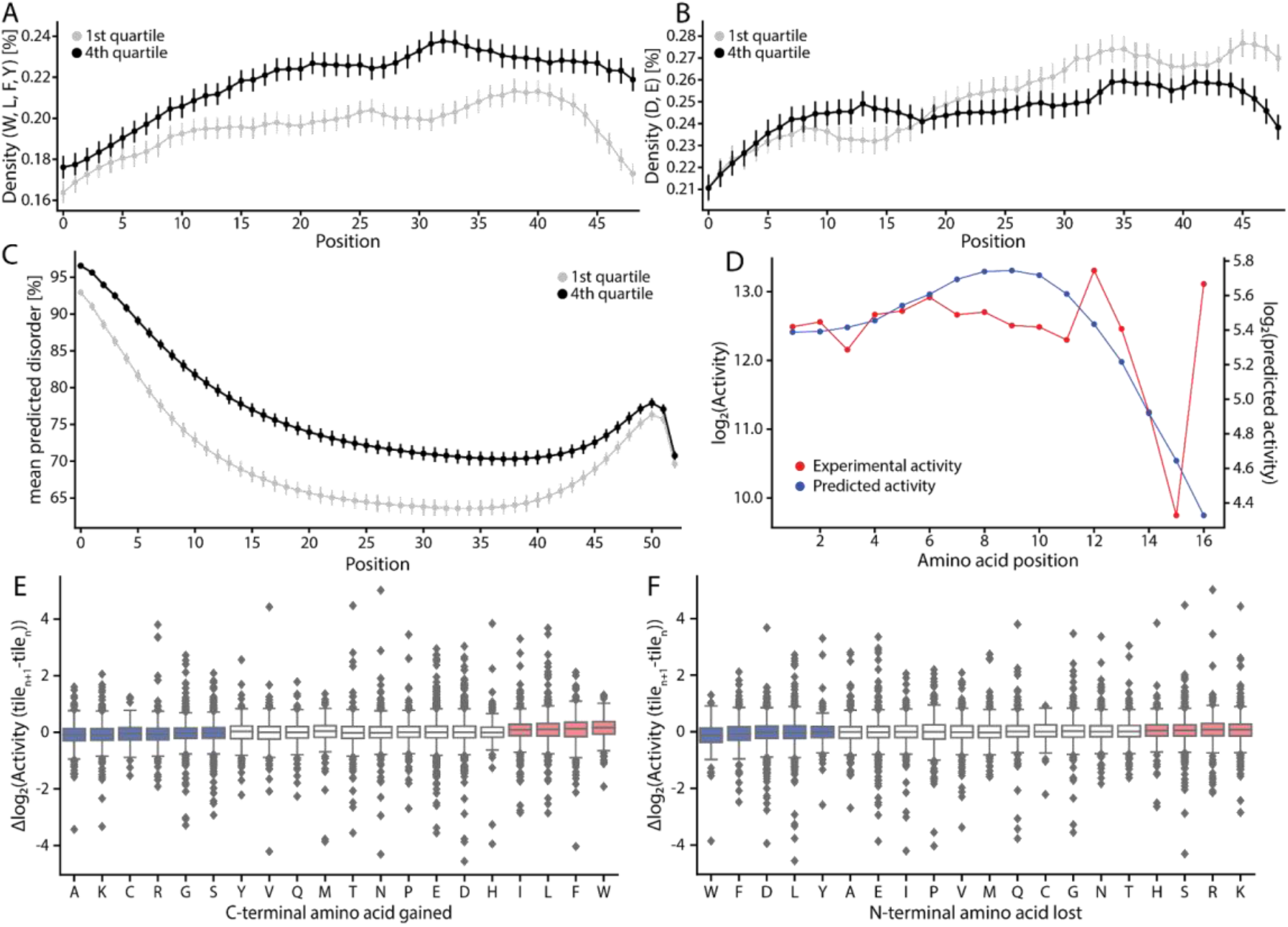
Tiling ADs with single amino acid resolution deciphers positional distribution of key amino acid groups. Density of functional amino acids across every position of every tile in the quartile with the strongest and weakest activity (4388 sequences per quartile). Density is calculated in a five amino acid window for each position along the AD as the average of all (**A**) hydrophobic residues (W, L, F, Y), (**B**) acidic residues (D, E). Error bars indicate the 95% confidence interval. (**C**) Mean predicted disorder at every position of tiles fused to the synthetic TF in respective quartiles predicted by MetapredictV2. Error bars indicate the 95% confidence interval. (**D**) Example of PADDLE predicted vs measured activity in single amino acid resolution of regions of interest (AT1G14630). Tiling protein sequences with single amino acid resolution allows us to observe the effects on activity when (**E**) gaining a C-terminal amino acid or (**F**) losing an N-terminal amino acid. Blue colored boxes indicate amino acids decreasing activity, red boxes indicate amino acids increasing activity.

We further studied the role of intrinsic disorder on activity in our library. Virtually all activation domains are intrinsically disordered, and the acidic exposure model suggests that more disordered sequences are likely more active. To gauge the disorder of tiles fused to synthetic TFs in the respective quartile, we utilized the disorder predictor Metapredict V2 (31). We found that all quartiles displayed increased disorder in their N-terminus, suggesting that initial disorder in the tile is important for activity **(Fig. 2C)**. In all quartiles, disorder dropped drastically in the C-terminus and the fourth quartile showed increased disorder throughout the entire tile **(Fig. S8)**. The disorder in the N-terminus implies that an entropic spacer or expanded linker between the estrogen binding domain and the AD increases activity. It is further possible that some sequences in this library may be interacting with the estrogen binding domain, which could drive collapse and decrease activity. The drop in predicted disorder at the C-terminus is likely the consequence of the increased density of aromatic residues (**Fig. 2C**). In consequence, current predictive models are still missing necessary positional information of key residues in ADs that need to be incorporated into future network training.

High-throughput studies usually scan protein sequences by tiling in step sizes of >10 amino acids (1, 8). Here we decided to tile at single amino acid resolution to study how single amino acid changes both from losing and gaining one amino acid during tiling affects AD activity **(Fig. 2D, SI Data 2)**. At the C-terminus, gaining the hydrophobic residues (W, F, L) enhanced activity as expected. Notably, isoleucine, which is not normally linked to enhancing activity, had a stronger positive effect on activity than acidic residues **(Fig. 2E)**. The enrichment of isoleucine was only observed in the C-terminal position tiles **(Fig. S9)**, suggesting an unknown role of this residue in AD activity. At the N-terminus all effects were smaller than for the C-terminus, but losing hydrophobic residues or aspartate decreased activity, and losing positively charged residues arginine and lysine increased activity following known rules of activity **(Fig. 2F)** (**6, 9**). Overall, we show that the amino acid composition alone cannot fully explain AD activity, with the position and distribution of key amino acids playing a critical role.

### A genome-wide compendium of coactivator candidates in plants

Coactivators provide an interface between TFs and RNA polymerase and are essential for the activation of gene expression. Although there has been significant attention on characterizing TFs at a genome-scale, only a limited number of coactivators have been characterized in plants, limiting our ability to fully understand how they regulate transcription. Artificially recruiting coactivators and chromatin regulators to DNA can directly modulate transcription (4, 32). Moreover, coactivators can contain ADs. We reasoned that ADs could occur in any gene involved in transcription because nuclear non-TF genes with ADs represent potential coactivators. Based on subcellular localization data, our library contains ADs from 107 and 200 nuclear non-TF genes in yeast and Arabidopsis, respectively, allowing us to explore their potential coactivator function **(Fig. 3A, B)**.

**Figure 3.**
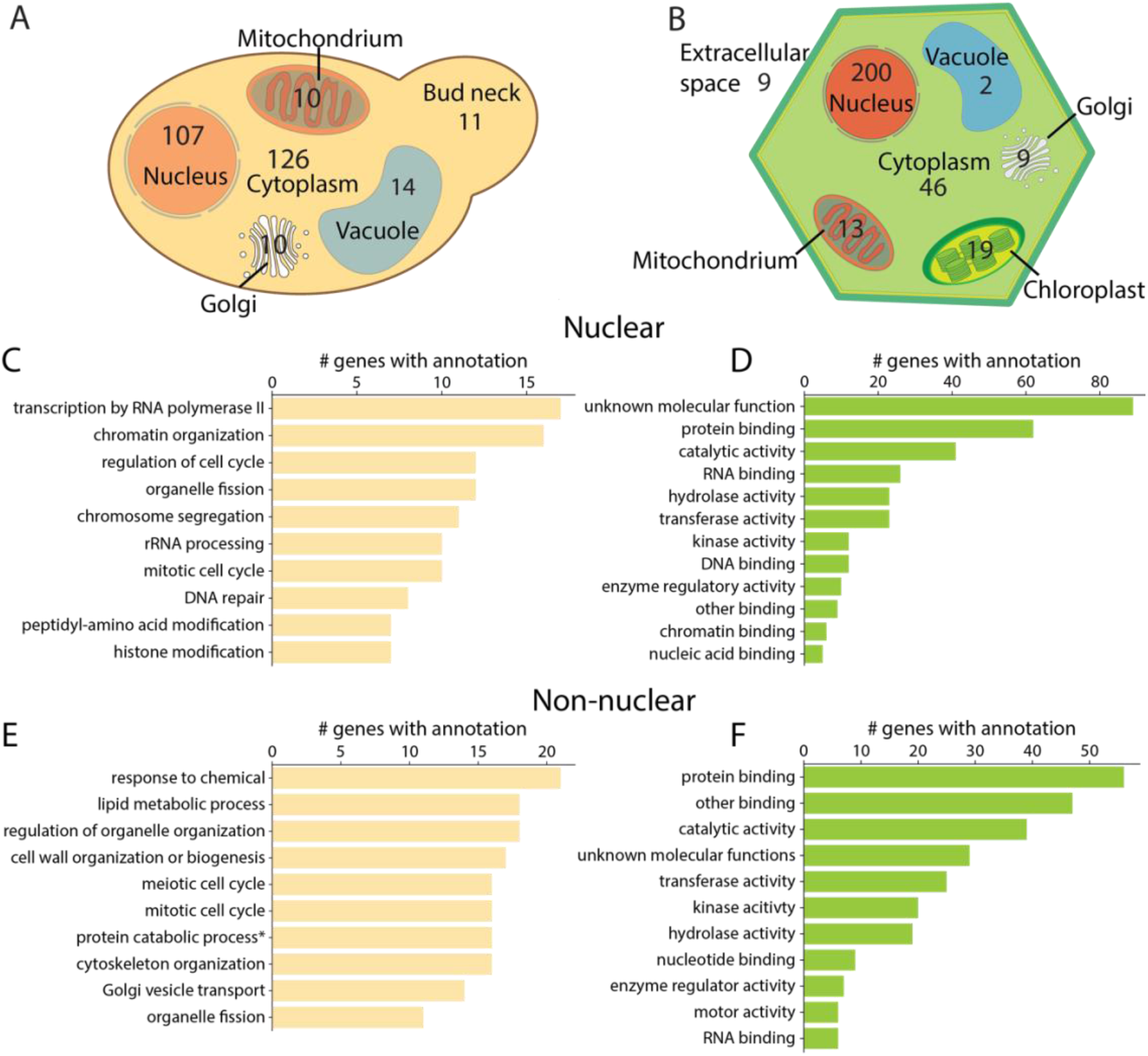
Non-nuclear proteins occurring throughout organelles have predicted ADs in both yeast and plants. Cell schematics with occurrence of parent genes with ADs in (**A**) *S. cerevisiae* and (**B**) *A. thaliana*. GO terms associated with nuclear genes in (**C**) *S. cerevisiae* and (**D**) *A. thaliana*. GO terms associated with non-nuclear genes in (**E**) *S. cerevisiae* and (**F**) *A. thaliana*.

Coactivators have been more thoroughly studied in yeast than in plants; hence, we benchmarked the occurrence of ADs in nuclear non-TF genes from yeast both to identify previously unannotated coactivator candidates as well as provide a more comprehensive list of known coactivators with ADs. To ensure only annotating genes with functional ADs, we included parent genes that yielded the 50% strongest ADs in the library. We used gene ontology (GO) terms to gauge the function of candidate genes and found most GO terms to be linked to transcription such as ‘transcription by RNA polymerase II’, ‘chromatin organization’, ‘regulation of cell cycle’ and ‘histone modification’ **(Fig. 3C, SI Table 5)**. As expected, we characterized tiles derived from known coactivators in yeast, namely IFH1, MED2, ROX3 and NRS1 (33–37). We also found ADs in chromatin regulators, namely HFI1, CHD1, SFH1, STH1, SDS3, CTI6 and INO80 (35, 38–43). Tiles from other proteins involved in transcription included transcription initiation factor eIF4G1, TATA binding factor TAF1, TAF14 and BDF1 as well as general TF TFG1 (44–47). Notably, we found ADs in two genes of unknown protein family and function (YBL029W, YML108W) which may function as potential co-activators. Candidate genes were also associated with the GO terms ‘rRNA processing’ and ‘chromosome segregation’, raising the question what roles ADs might play in these proteins. Overall, previous observations of ADs in coactivator complexes and chromatin regulators were supported by our results, our approach can be used to annotate putative genes with regulatory function.

Coactivators in plants have been far less studied and mostly annotated based on homologs from other eukaryotes (48, 49). Hence, the parent genes of tiles from Arabidopsis contained far fewer hits in known transcription associated genes. Of the 211 nuclear non-TF hits, only 4 had previously been validated to be coactivators, highlighting the opportunity to discover new plant coactivators. We found ADs in the coactivators MED13 and LNK1/LNK2/LNK3 (50–52), the chromatin regulators HAF2 and SCS2A/B (53, 54), and four transcription elongation factors from family S-II. We also found ADs in three members of the VQ family of suspected transcriptional coregulators that interact with WRKY family TFs during abiotic stress response and four CCT-motif-containing proteins that have been linked to transcriptional elongation in other eukaryotes (55). Notably, only 23 genes have GO terms linked to transcription with terms like ‘chromatin binding’, ‘nucleic acid binding’, ‘DNA binding’. The most abundant GO term was ‘unknown molecular functions’ with 89 associated genes, highlighting putative coactivators that have not yet been characterized **(Fig. 3D, SI Table 6)**. Other nuclear genes with ADs were either not previously associated with transcription or have never been studied before, suggesting that there may be plant-specific coactivators that cannot be annotated purely based on sequence homology to other eukaryotes. Our results supply an extensive list of putative coactivators in Arabidopsis which should accelerate the proper characterization of such proteins; ultimately providing useful targets to modulate and engineer key traits in plants.

### Non-nuclear proteins throughout all organelles contain ADs

Non-nuclear proteins can contain ADs and influence transcription via relocation to the nucleus as exemplified in the examples of Notch1 and beta-catenin (11, 12). We investigated the prevalence of ADs that occur in proteins outside of the nucleus. We again only focused on parent genes harboring tiles of the 50% strongest experimentally validated ADs, yielding 136 Arabidopsis and 207 yeast non-nuclear genes **(Fig. 3A, B)**. We found 46 and 126 cytosolic in Arabidopsis and yeast, respectively. These genes are candidates for relocalization into the nucleus, similar to Notch1 and beta-catenin. Besides these candidates in Arabidopsis, we found AD containing genes in the chloroplast (19), plasma membrane (15), mitochondria (13), golgi (9), endoplasmic reticulum (9), peroxisome (2), vacuole (2) and extracellular space (9). In yeast there were candidates in the endoplasmic reticulum (12), vacuole (14), bud neck (11), mitochondria (10), the vacuolar membrane (3) and multiple non-nuclear organelles (26). Overall, 90 Arabidopsis and 101 yeast non-nuclear genes with ADs are non-cytosolic and are targeted to a specific organellar compartment, raising the question whether they can be relocated to the nucleus to modulate transcription.

To gauge the role of all non-nuclear candidate genes, we studied their GO terms in both species. In yeast, GO terms were unrelated to transcription and included both metabolic terms like ‘lipid metabolic process’, and signaling terms like ‘response to chemical’. Many GO terms were linked to architecture of the cell like ‘meiotic/mitotic cell cycle’, ‘cytoskeleton organization’ and ‘organelle fission’ **(Fig. 3E, SI Table 7)**. In Arabidopsis, the two most abundant molecular function terms were linked to “protein binding” and “general binding”, followed by “catalytic activity” and “unknown molecular function” **(Fig. 3F, SI Table 8)**. The inclusion towards GO terms linked to binding interactions in Arabidopsis raises the question whether AD-like peptides have been co-opted in other organelles to facilitate protein-protein interactions outside of the nucleus. In principle, isolating proteins in topologically separate compartments enables the reuse of the same protein-protein interactions without cross-talk. This highlights the diverse functionalities of proteins with AD-like sections in both species and suggests that AD-like peptides may perform other protein-protein interactions outside the nucleus.

### A set of universal eukaryotic activation domains

Our library gave us the unique opportunity to validate the transferability of yeast derived ADs into plants and establish potential universal activators that function in phylogenetically divergent eukaryotes. We characterized 55 of the strongest ADs in the library – 33 derived from Arabidopsis and 22 from yeast – in the plant *Nicotiana benthamiana* using an agroinfiltration-mediated transient expression system that we previously established **(SI Table 9)** (10). In this system each tile is fused to the yeast GAL4-DBD and localized to a synthetic minimal promoter with 5 concatenated GAL4 binding sites to drive GFP **(Fig. 4A)**, modulating GFP expression. A constitutively expressed dsRed is used to normalize the signal. To benchmark the potency of the AD tiles we also tested the strong activator VP16 and VPR (a fusion of the three strong activators VP64, p65 and RTA) as GAL4 fusions. Of 55 ADs tested, we found 43 (78.2%) to significantly increase GFP expression over the reporter only control, 6 were stronger than VP16 and 2 were statistically indistinguishable from VPR (Mann-Whitney-U test, p < 0.05) **(Fig. 4A, SI Table 10)**. Notably, our tested ADs span the entire dynamic range of possible activities *in planta*, from no observed to very strong activity, highlighting the importance of remeasuring ADs *in planta*. Overall, we discovered short, universal ADs from non-TF proteins that perform similarly to longer state of the art ADs like VPR that are readily available for further eukaryotic engineering efforts. Shorter and stronger ADs are in demand for space-limited synthetic biology engineering strategies such as viral delivery methods.

**Figure 4.**
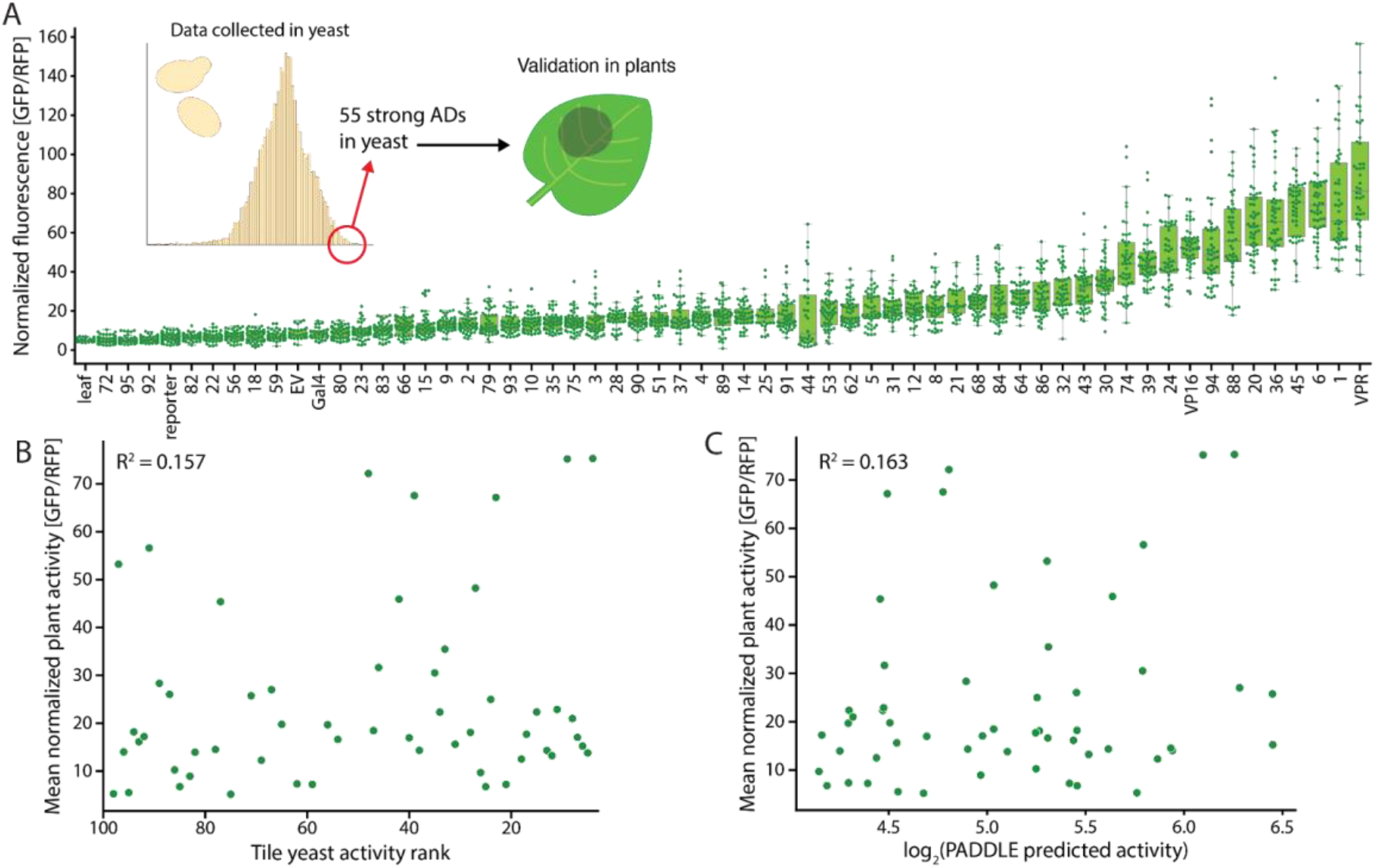
Cross-validation of strong ADs yields candidates with similar activities to gold standard activators. (**A**) Activity of 55 of the 100 strongest tiles measured in yeast in *N. benthamiana*. VP16 and VPR serve as positive AD controls. EV: Empty Vector control, Gal4: Gal4-DBD without any AD. (**B**) Mean normalized *in planta* activity of ADs in (A) sorted by activity strength in yeast. (**C**) Mean normalized *in planta* activity of ADs in (A) relative to predicted activity.

Our agnostic approach to mapping ADs in non-TF genes allowed us to mine strong ADs from proteins that have not previously been associated with transcription and we show that these ADs function in plants. As an example, we localized and validated ADs in known plant coactivators, namely one CCT-motif containing protein from Arabidopsis (AT1G04500), coactivator LNK3 (AT3G12320) and SAGA complex subunit 2A (AT2G19390), which showed activity similar to the VP16 control. Furthermore, the strongest AD in plants was derived from an Arabidopsis uncharacterized 2Fe-2S ferredoxin-like superfamily protein (AT1G50780). The second strongest AD was derived from a hypothetical protein (AT2G29920). Overall, ADs in non-TF proteins involved in transcription and in non-nuclear proteins function *in planta*.

Eukaryotic TFs utilize conserved general transcription machinery (e.g., Mediator) to facilitate transcription, making new TF parts a potential resource to develop tools for the control of transcription across eukaryotes. For our plant experiments we chose only the strongest tiles from our yeast experiments, expecting that, if they utilize general conserved transcription machinery, activities in plants should be similarly strong. However, we observed poor correlation between the rank order of yeast AD versus the rank order in plants **(Fig. 4B)**. PADDLE predictions correlated worse with observed AD activity in plants than in yeast **(Fig. 4C)**. Our results suggest that, while PADDLE can localize ADs on parent genes in both plant and yeast proteins with 89.0% accuracy, there are mechanistic features of plant transcription not fully captured by PADDLE that prevent accurate prediction of AD strength. We conclude that future work is needed to generate independent plant AD datasets to train models that can predict the strength of plant ADs with higher accuracy to enable the full potential of mining plant proteomes.

## Discussion

High-throughput studies have largely focused on ADs found in TFs and protein classes known to be involved in transcription, which has partly biased our understanding of the biological role of such peptides. By mining proteomes for ADs from non-TF genes and demonstrating their activity in yeast and plants, we reveal that ADs frequently occur across entire proteomes and outside the nucleus, going beyond the canonical description of ADs in TFs that mediate nuclear transcription. Studying nuclear non-TF ADs from the well-studied model yeast expands our understanding of which genes contain AD-like peptides and where they are localized. We found a direct correlation between nuclear genes containing ADs and their likelihood to function as coactivators in yeast. Our dataset provided the motivation to extrapolate this observation to plants and we annotated over 200 putative coactivators that may be involved in many facets of plant transcriptional regulation. Due to the throughput limitations of our experimental setup, we focused on the strongest 18,000 tiles from both species, leaving a far larger sequence space of medium or weak ADs unstudied. Future work will focus on experimentally validating larger sets of predicted ADs in both species and help understand how frequent ADs occur throughout proteomes.

The recent establishment of large experimental datasets of activation domains in yeast has led to the development of multiple neural networks that attempt to localize and predict the activity of ADs from protein sequences (1, 6). In this study we utilized one of these models PADDLE to build and test our library (1). We found that PADDLE can correctly localize ADs throughout entire proteomes; however, the capabilities of PADDLE to predict the quantitative activity of ADs fell short in comparison to the high correlation value that was reported in the original study. We further show that plant tiles that functioned as strong ADs in yeast, indeed largely functioned in plants but with divergent degrees of activity. This discrepancy indicates that while general eukaryotic mechanisms for the regulation of transcription between plants and yeast are conserved, there are intricacies in plants that models trained on yeast data cannot resolve. These intricacies parallel leucine-dependent ADs from metazoans that are missing in yeast (15). It further highlights that the flexible positional and compositional sequence requirements of ADs need to be explored further in their native context.

Recently, there has been significant interest in utilizing genome engineering approaches in non-model eukaryotes that have traditionally been recalcitrant to genetic studies. Such efforts are constrained by the dearth of characterized genetic parts that reliably function across phylogenetically diverse eukaryotes, restraining the application of high-throughput genetic screening methods, like CRISPR activation (CRIPSRa). Our results highlight how computational models for predicting ADs are still in their infancy. Sequences with very similar amino acid composition can largely differ in activity based on amino acid arrangement. Future work is needed to further unravel this sequence “grammar” to better understand AD function and guide the construction of next-generation predictive models. In the long term, we anticipate building models for species-specific activity. Such knowledge will help establish design principles for ADs which will ease the implementation of new synthetic biology tools and genome-scale activation assays like CRISPRa screens in plants and other non-model organisms.

At their core, ADs enable protein-protein interactions with transcriptional machinery to facilitate transcription. We observed abundance of peptides with AD-like properties throughout entire proteomes in two distantly related eukaryotes, suggesting three possible roles of these peptides. 1) Their parent proteins moonlight into the nucleus to facilitate transcription. 2) The broad sequence space that allows AD activity has lead to statistical occurrence of peptides with AD-like properties. 3) ADs are an instance of a larger class of protein-protein interaction domains that perform many functions. Option one is supported by anecdotal evidence of proteins normally localized to the cytosol and mitochondria being imported into the nucleus to promote transcription during signaling cascades (11, 12, 56). Option two is supported by large AD screens that show that up to 1% of random sequences have AD activity when localized to promoters (57–59). We believe that option three entails the most logic. ADs do not rely on structure to facilitate binding, they form multiple weak interactions with coactivators (60, 61). We believe that compartmentalization allows the ‘recycling’ of AD-like interactions in different organelles. Of note, the gene ontologies predicted from physically separated plant organelles (e.g., mitochondria and chloroplast) are highly enriched in proteins involved in “protein binding,” supporting that these interactions could be used for different functions independent of transcription. We speculate that the versatile nature of ADs extends their role beyond nuclear transcription and blurs the distinction of ADs as a feature unique to transcription factors.

## Material and methods

### Data sharing plans

Code - We will generate a timestamped GitHub repository of all code used for analysis after review

Sequencing - We will deposit all sequencing raw reads used in this study on GEO

Data - All other data is available as SI

### PADDLE prediction of every 53 amino acid tile in the proteome of A. thaliana and S. cerevisiae

We predicted the AD activity of all proteins of the reference proteome of *A. thaliana* (Colombia ecotype) and *S. Cerevisiae* (strain S288C) which we obtained from TAIR (Araport11) and SGD (S288C Genome release 64-3-1), respectively. Both proteomes with associated predictions are available in Supplementary data file 1 and can be loaded using Load_predictions_SI_data1.ipynb. We predicted the secondary structure of every full-length protein using S4PRED and their structural disorder with IUPRED3 (long and short mode) (62, 63). We then tiled the protein sequences and structural predictions into consecutive 53 amino acid tiles and predicted their AD activity using the PADDLE API for Python as described (1). We ran all predictions in Python v3.9.5 with associated APIs and our pipeline is available in the Supplementary Data package. As we wanted to focus on tiles from non-TF genes, we utilized the TF databases PlantTFDB v5.0 and Yeastract+ to filter out any tiles derived from TFs. We selected tiles from genes that achieved a PADDLE predicted activation >30, yielding 12,000 *A. thaliana* tiles and 6,000 *S. cerevisiae* tiles with a dynamic range of PADDLE predicted activation strength between 17 and 138.

### Plasmid library construction

The library of both Arabidopsis and Saccharomyces ADs were generated by mapping the tiles back to their native DNA sequence in the respective reference genomes, retrieved from TAIR and SGD (all sequences in SI Table 1). 18,000 unique DNA oligos coding for 53 amino acid long putative activators were synthesized in one oligo pool by Twist Bioscience. Each oligo contains a 24 bp upstream primer (GCGGGCTCTACTTCATCGGCTAGC), 159 bp encoding the activator candidate, a 21 bp primer (TGATAACTAGCTGAGGGCCCG) with four stop codons in 3 frames and the ApaI site. Specifically, we used 75 ng of template and 12 rounds of PCR in 16 parallel 50 μL reactions using primers LC3.P1_Lib_Hom_up_5’, which adds homology arms and YL_randBCs_R1_3’ which adds random 11 nt barcodes and downstream homology arms (NEB Q5 polymerase Tm=70C). The PCR product was pooled and cleaned using the Monarch PCR and DNA kit, followed by product visualization on a 1% Agarose gel. Vector pMVS219 was linearized using NheI, AscI and PacI and used for library assembly. The assembly was performed using 100 ng of linearized backbone and 7.5 ng of PCR product using NEB Hifi DNA Assembly Master Mix in 8, 10 μL reactions. Assemblies were electroporated into DH5β cells (NEB C3020K), and we recovered >1,000,000 colonies.

The plasmid sequence of the library assembly vector pMVS219 is available on addgene (https://www.addgene.org/99049/).

Primers:

YL_randBCs_R1_3’ AATTCGCTTATTTAGAAGTGGCGCGCCNNNNNNNNNNNCGGGCCCTCAGCTAGTTATCA

LC3.P1_Lib_Hom_up_5’ GCGGGCTCTACTTCATCGGCTAGC

### Yeast Library Construction and measurement

To ensure singular constructs per cell, we introduced our library into the URA3 locus of strain DHY211 (*MATa, MKT1(30G,) RME1(INS-308A) TAO3(1493Q),CAT5(91M) MIP1(661T), SAL1^+^ HAP1^+^*). Employing the established yeast transformation method (64), we subjected the transformation to 30 minutes at 30 °C followed by 60 minutes at 42 °C. To minimize potential PCR errors, we performed SalI and EcoRI digestion on the plasmid library, releasing the section encompassing the ACT1 promoter, the synthetic TF, and the KANMX marker. Simultaneously, PacI digestion was conducted to cleave plasmids devoid of an activation domain variant and barcode insert, thereby reducing the occurrence of transformants with inactive TFs. Directed integration into the URA3 locus was guided by 500 bp upstream homology spanning the URA3 and ACT1 promoters, along with a corresponding 500 bp downstream homology region spanning the TEF and URA3 terminators. These regions were PCR amplified from pMVS 295 (Strader 6161) and pMVS 296 (Strader 6768), a generous gift from Nick Moffy and Lucia Starder. Transformation utilized a molar ratio of 1:3 for linearized library to homology arms, with 28 μmol of linearized library per reaction. The transformed library was plated on YPD, followed by an overnight incubation at 30°C, and subsequent replica-plating onto freshly prepared SC G418 plates. Employing this process across 80 transformation reactions yielded an estimated >1,000,000 individual colonies. Subsequently, the transformants were collectively mated with an FY5 strain containing the reporter integrated into the uncertain ORF, YBR032w. Diploids were selected on YPD with G418 (200 µg/ml) and NAT (100 µg/ml) (strain MY436 YBR032w::P3_GFP NAT S288C), resulting in prototrophic diploids. These 110,000 yeast transformants were mated in batches, and prior to the final experiment, batches were pooled and multiple aliquots were frozen.

### Fluorescence Activated Cell Sorting and library preparation

Each sorting experiment was preceded by thawing a frozen glycerol stock, followed by overnight growth in SC+G418+NAT. Cultures were cultivated in synthetic complete (SC) dextrose media at 30 °C (65). Prior to fluorescence-activated cell sorting (FACS), overnight cultures were diluted (1:5) into SC+ 1 μM ß-estradiol and incubated for 3.5-4 hours at 30 °C. We sorted the yeast library on a Aria-fusion cell sorter at the UC Berkeley Flow Cytometry core facility. We used the parent yeast strain with the reporter and a TF lacking an activation domain as a negative control to determine autofluorescence and baseline mCherry levels. We sorted 1 million cells of the synthetic TF library into 8 bins with each bin roughly covering 11 % of the entire observable population in the GFP channel. To test reproducibility, we sorted another 500,000 cells from each bin.

Sorted cells were grown overnight in SC at 30 °C and gDNA was extracted with the Zymo YeaSTAR (#D2002) kit. Barcodes were amplified by PCR (CP21.P14: TCCTCATCCTCTCCCACATC, CP17.P12: GGACGAGGCAAGCTAAACAG, NEB Q5 for 20 cycles, Tm 67 °C). We added phasing nucleotides as well as overhangs for indexing primers using primer mixtures SL5.F[1-4] and SL5.R[1-4] (NEB Q5 for 20 cycles, Tm 62 °C). We finally added dual indexing primers using the i5 and i7 system from Illumina (NEB Q5 for 20 cycles, Tm 65 °C). We then performed a bead cleanup. We sequenced the library on an Illumina Novaseq 6000 system with 2×150 bp paired end reads.

We assessed library performance against known ADs from GCN4 and VP16 on a BD Accuri^TM^ C6 flow cytometer (BD Biosciences). All strains were grown in SC+G418+NAT at 30 overnight and diluted (1:5) into SC+/- 1 μM ß-estradiol and incubated for 3.5-4 hours at 30 °C. Samples were washed with cold 1x PBS (137 mmol NaCl, 2.7 mM KCl, 1.8 mM KH2PO4, 10 mM Na2HPO4) once before measurement. Per sample 100,000 events were recorded and analyzed using the Python fcsparser package.

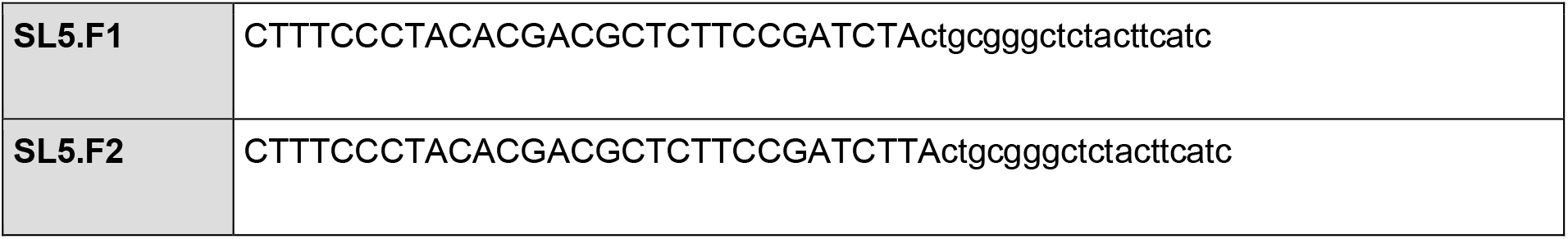
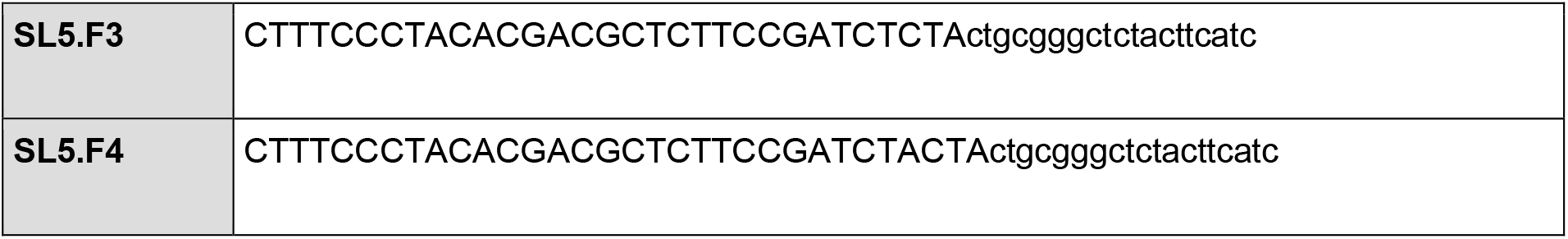
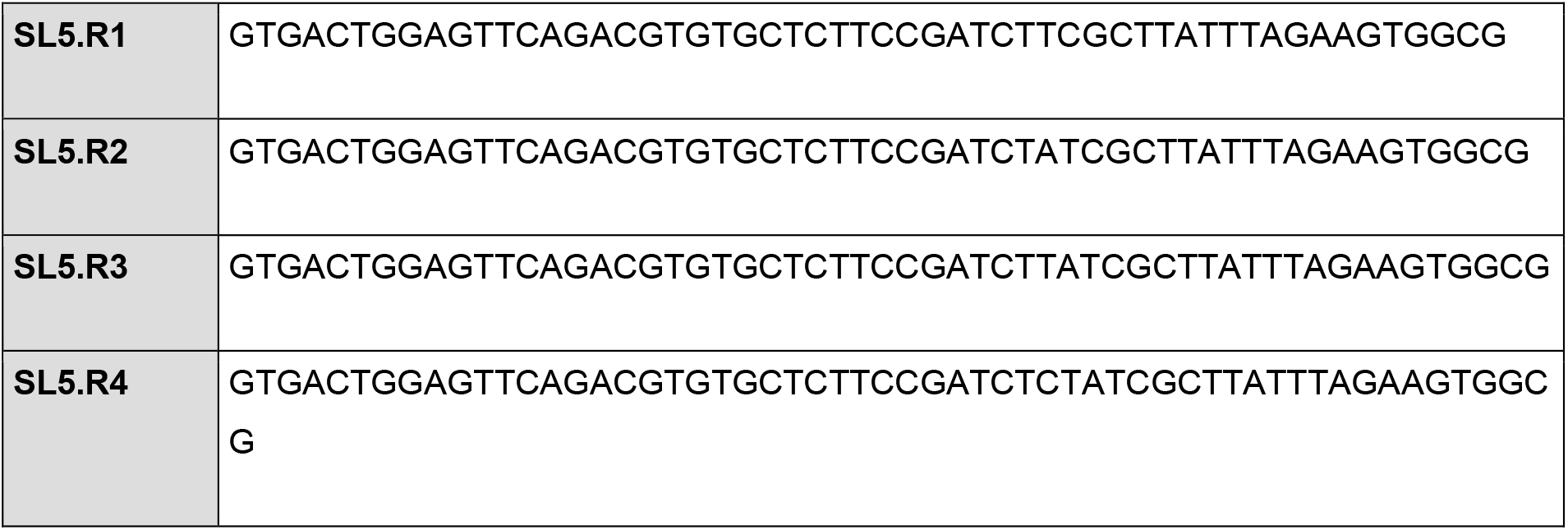

### Plant experiments

Generated binary vectors were transformed into *Agrobacterium tumefaciens* strain GV3101. Selected transformants were inoculated in liquid media with appropriate selection the night before the experiment. *A. tumefaciens* strains were grown until OD600 between 0.8 and 1.2 and were mixed equally (final OD600 = 0.5 for each strain) with the strain harboring the assay reporter construct to a final OD600 = 1.0. Cultures were centrifuged for 10 min at 4000 g and resuspended in infiltration buffer (10 mM MgCl2, 10 mM MES, and 200 μM acetosyringone, pH 5.6). Cultures were induced for 2 h at room temperature on a rocking shaker. Leaves 6 and 7 of 4 week old *N. benthamiana* plants were syringe infiltrated with the *A. tumefaciens* suspensions. Post infiltration *N. benthamiana* plants were maintained in the same growth conditions as described above. Leaves were harvested three days post infiltration and 16 leaf disks from two leaves and 3 plants total per construct were collected. The leaf disks were floated on 200 µL of water in 96 well microtiter plates and GFP (Ex. λ = 488 nm, Em. λ = 520 nm) and RFP (Ex. λ = 532 nm, Em. λ = 580 nm) fluorescence measured using a Synergy 4 microplate reader (Bio-tek). The reporter construct for the screen was pms6370 containing GFP and dsRed expression cassettes. GFP expression was driven by a fusion of five previously characterized GAL4 binding sites with the core WUSCHEL promoter (66). GFP expression was normalized using dsRed driven by the constitutive MAS promoter on the same plasmid.

### Analysis of barcodes and inferring activity

After demultiplexing samples, we kept only the reads that contained a perfect match to a designed tile. For each set of 8 sorted samples, we performed two normalizations. We first normalized the reads by the total number of reads in each bin. Then, for each designed tile, we normalized across the 8 bins to calculate a relative abundance. We then converted relative abundances to an activity score for each tile by taking the dot product of the relative abundance with the median fluorescence value of each bin **(SI Table 11)**. This computation is a weighted average. Tiles with less than 10 reads were not included in the final dataset. Later, post hoc analysis suggested that tiles with at least 1000 reads were well measured.

During plasmid library construction we added random barcodes to the designed tiles. To build a map linking designed tiles to barcodes, we combined all of the sequencing data from the 16 sorted samples. We use this map to compare two modes of analysis. First, for the primary analysis used in the manuscript, we used only the tile sequences, effectively combining all the barcodes together and ignoring independent transformations. Second, we repeated the analysis for each AD+barcode combination, in effect measuring the activity of each independent transformant of each tile. The methods largely agreed **(Figure S10)**. We determined statistical significance thresholds to infer the amount of tiles with AD activity. We calculated the statistical difference between each individual tile with the mean of no-AD control using a one sample t-test and corrected p-values using Benjamini-Hochberg false discovery rates (5%).

## Data analysis

We analyzed and visualized the data and underlying sequences of the tiles using the following APIs in Python v3.9.5: pandas, seaborn, matplotlib, numpy and scipy. All associated Jupyter Notebooks for producing all Figures are in the Supplementary Data files. We sorted the library by activity and split it into four equal sized quartiles with 4388 tiles per quartile. To gauge the composition of each tile in each quartile, we calculated the amino acid frequencies of all amino acids in each tile. For the amino acid density analysis we applied a sliding window size 5 along every position of each tile, averaging the frequencies of amino acid occurrence of each aminoacid for each quartile. We chose the amino acid window size to be 5 to not bias the analysis for short AD motifs like the 9aaTAD (67). We then grouped the amino acid frequencies based on functional groups which we defined as follows: acidic (D, E) and hydrophobic (W, L, F, Y).

To gauge the disorder of tiles we utilized the disorder predictor MetapredictV2 which integrated the outcomes of multiple independent disorder predictors (68). We predicted disorder of tiles when fused to the synthetic TF and in their endogenous context. Confidence intervals were calculated using the seaborn pointplot function.

Dipeptide frequencies were calculated by splitting tiles into quartiles as described before. We calculated the total occurrence of every amino acid in the respective quartiles. We measured the frequency of every dipeptide upstream and downstream, meaning if the first amino acid is an alanine, we accounted for all XA and AX dipeptides, where X is any of 20 twenty amino acids. The total occurrence of dipeptides was then normalized to the occurrence of the first amino acid in the quartile. We calculated dipeptide frequencies with spacers of up to 8 amino acids between amino acid one and two.

We provide figures of all parent genes with annotated location of tiles with their respective predicted and experimental activity as a resource in Supplementary data 3.

We utilized the single amino acid resolution of our tiling experiments to gauge the effect on AD activity when one C-terminal amino acid is gained or one N-terminal amino acid is lost. We generated a subset of tiles only including tiles that had at least one consecutive neighboring tile, meaning a pair of identical tiles with only one amino acid difference in the C- and N-terminus. From this subset we calculated the change of AD activity between consecutive pairs of tiles and associated the lost and gained amino acid during the step. The analysis was performed for the entire library independent of whether a tile was defined as an AD or not.

All code used for data analysis and associated data files are available on GitHub (https://github.com/shih-lab).

### Mapping putative coactivators

To generate a map of putative coactivators in both plants and yeast we firstly subdivided the library into genes with and without tiles with AD activity. Hence, we only studied parent genes of tiles in the 50% of the library that had higher activity. Then we further subgrouped parent genes into nuclear-genes by utilizing SUBA5 for Arabidopsis and YeastGFP/YPL+. To gauge their function we used Gene Ontology. For yeast we used the SGD GO term slim mapper and for Arabidopsis the functional categorization of GO terms in the bulk data retrieval tool. We then manually studied the molecular functions linked to all nuclear genes in both species to find known coactivators and chromatin regulators. We then excluded all known coactivators from our list to generate the final map of putative coactivators. For non-nuclear genes we used the same approach.

## Funding

This work was part of the DOE Joint BioEnergy Institute (http://www.jbei.org) supported by the U. S. Department of Energy, Office of Science, Office of Biological and Environmental Research through contract DE-AC02-05CH11231 between Lawrence Berkeley National Laboratory and the U.S. Department of Energy. The United States Government retains and the publisher, by accepting the article for publication, acknowledges that the United States Government retains a non-exclusive, paid-up, irrevocable, worldwide license to publish or reproduce the published form of this manuscript, or allow others to do so, for United States Government purposes. MVS is supported by NIH grant R35GM150813, Simons Foundation grant 1018719, and NSF grant 2112057. MVS is a Chan Zuckerberg Biohub – San Francisco Investigator.

---

**Figure S1.**
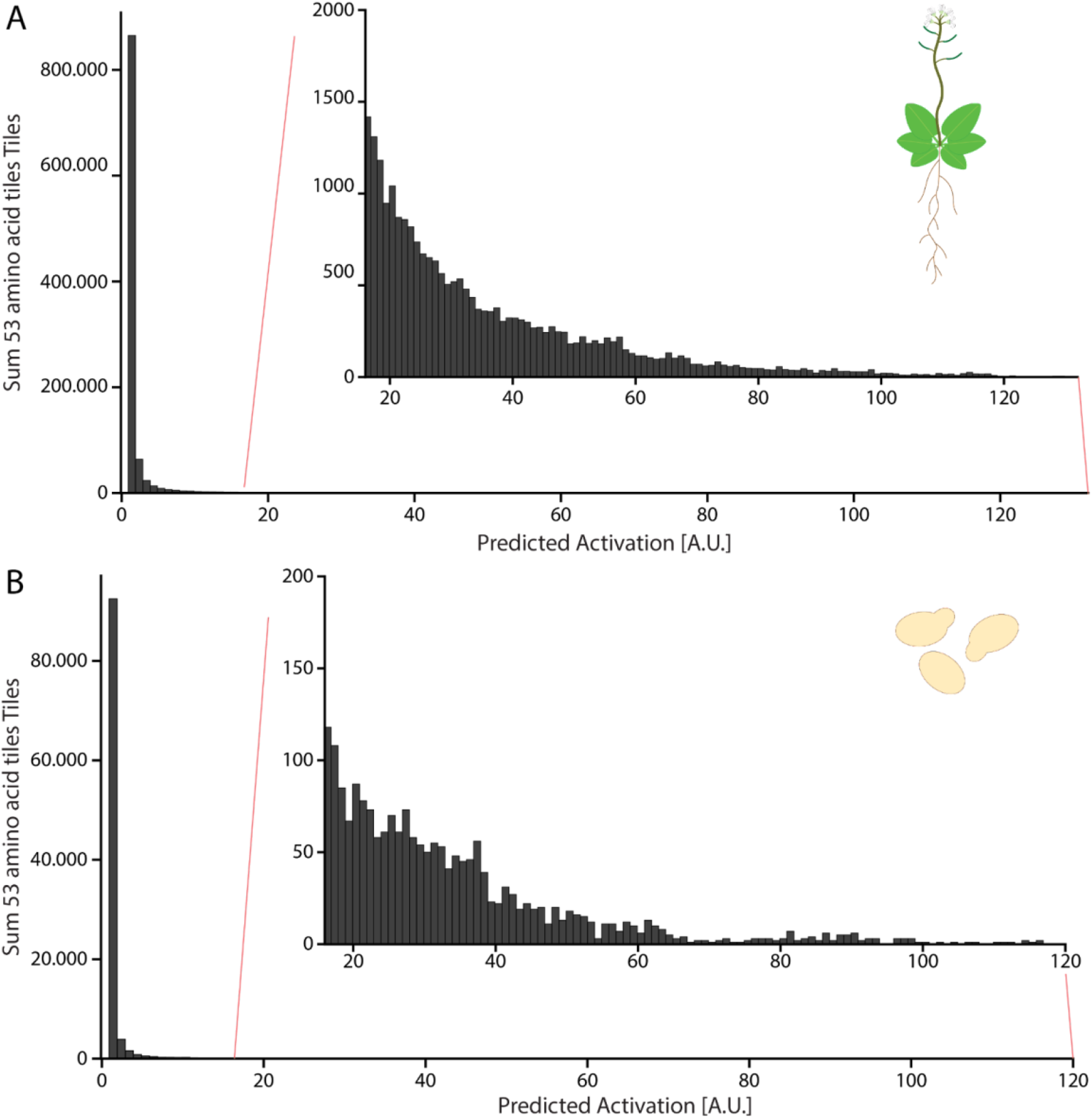
TFs in both Arabidopsis and yeast show similar predicted AD strength as non-TF genes. PADDLE predicted activation of every 53 amino acid tiles from all annotated (**A**) *A. thaliana* and (**B**) *S. cerevisiae* TFs. Inlets show the same area of predicted activity as used for choosing tiles for the library in Fig. 1A, B.

**Figure S2.**
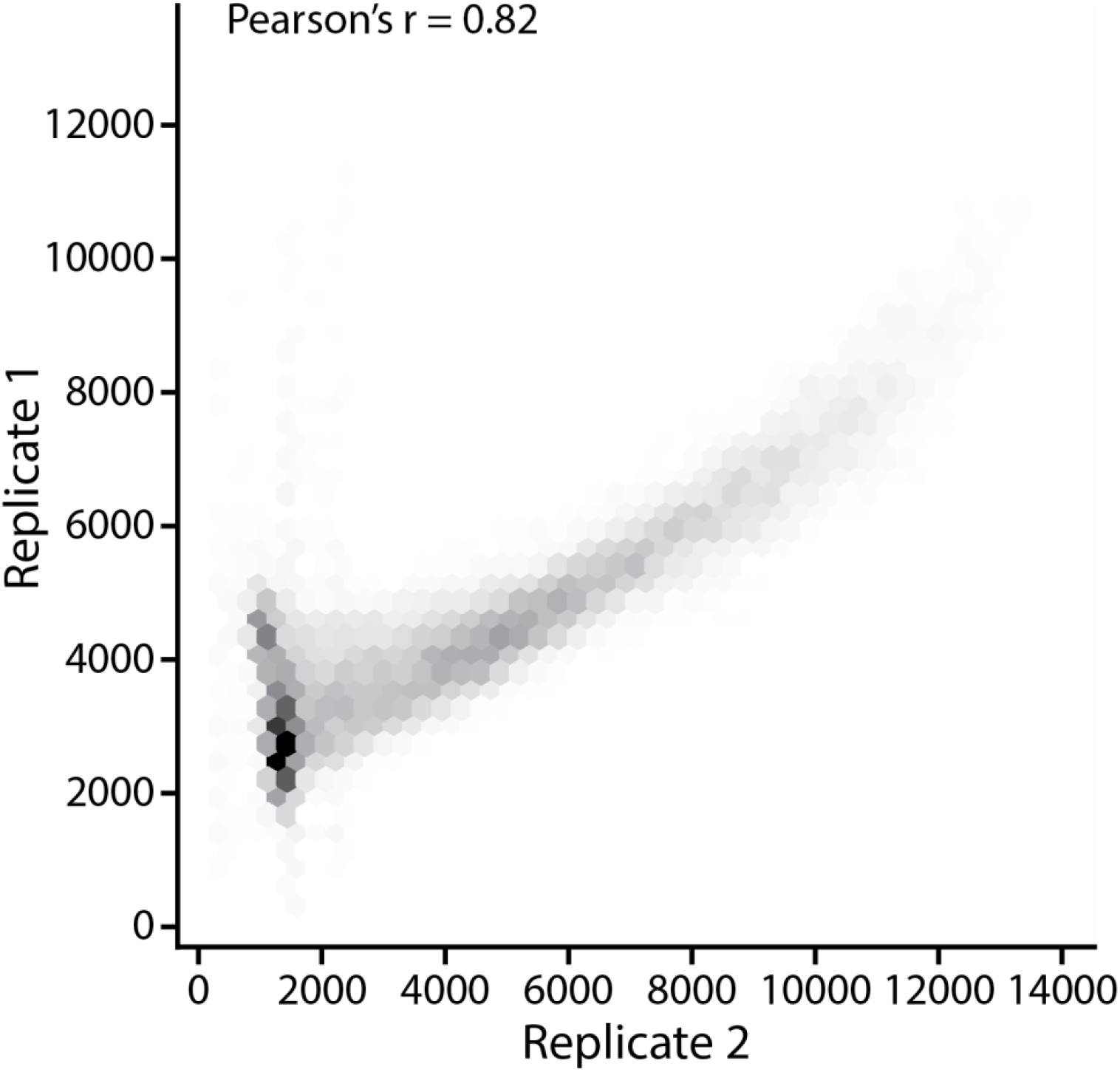
Activity of tiles is highly reproducible. Activities calculated from two independent sorts are strongly positively correlated.

**Figure S3.**
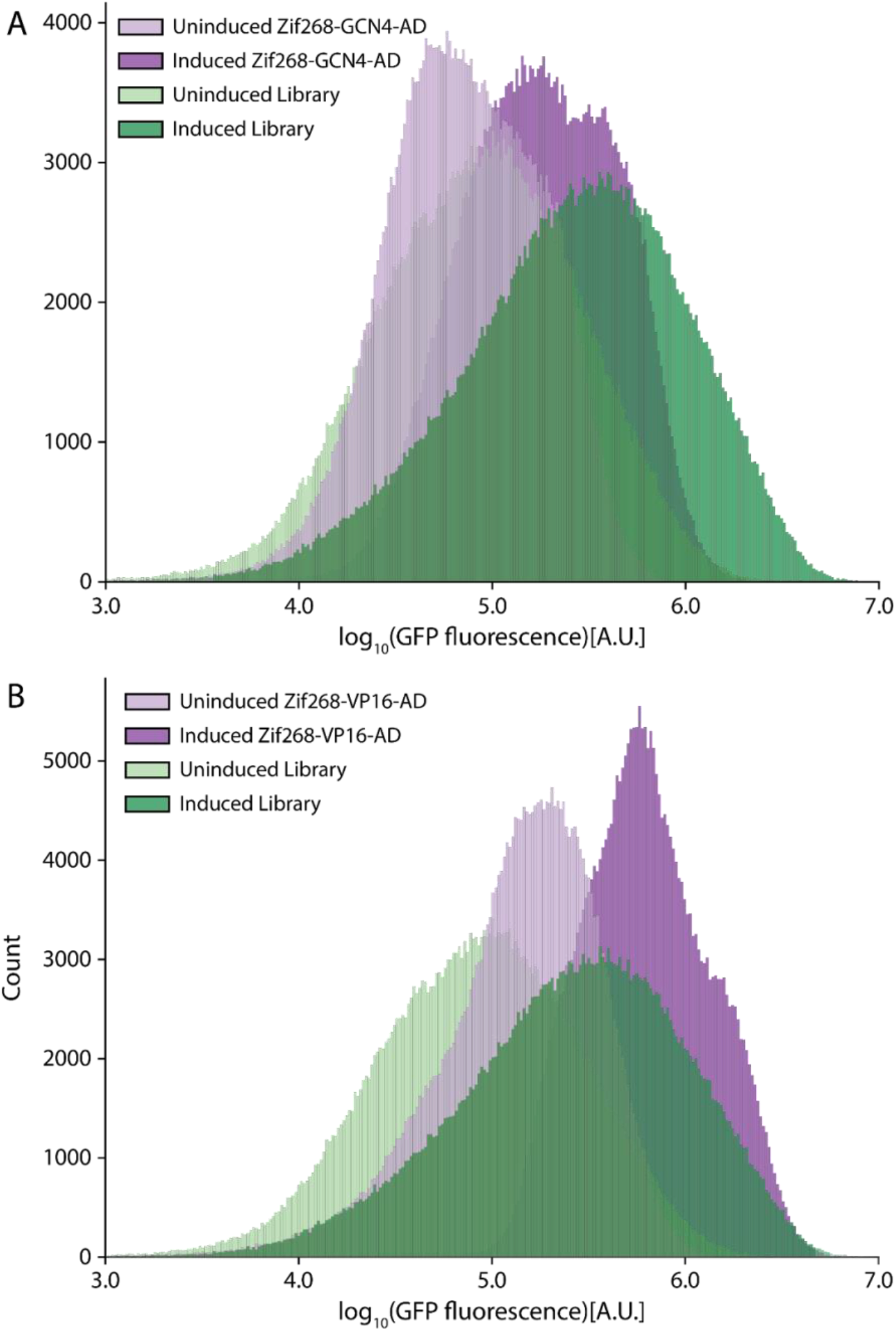
Flow cytometry of library against GAL4 and VP16 controls. Flow Cytometry comparing 100,000 observed events of the uninduced/induced library vs. (**A**) Zif268-GCN4-AD and (B) Zif268-VP16-AD.

**Figure S4.**
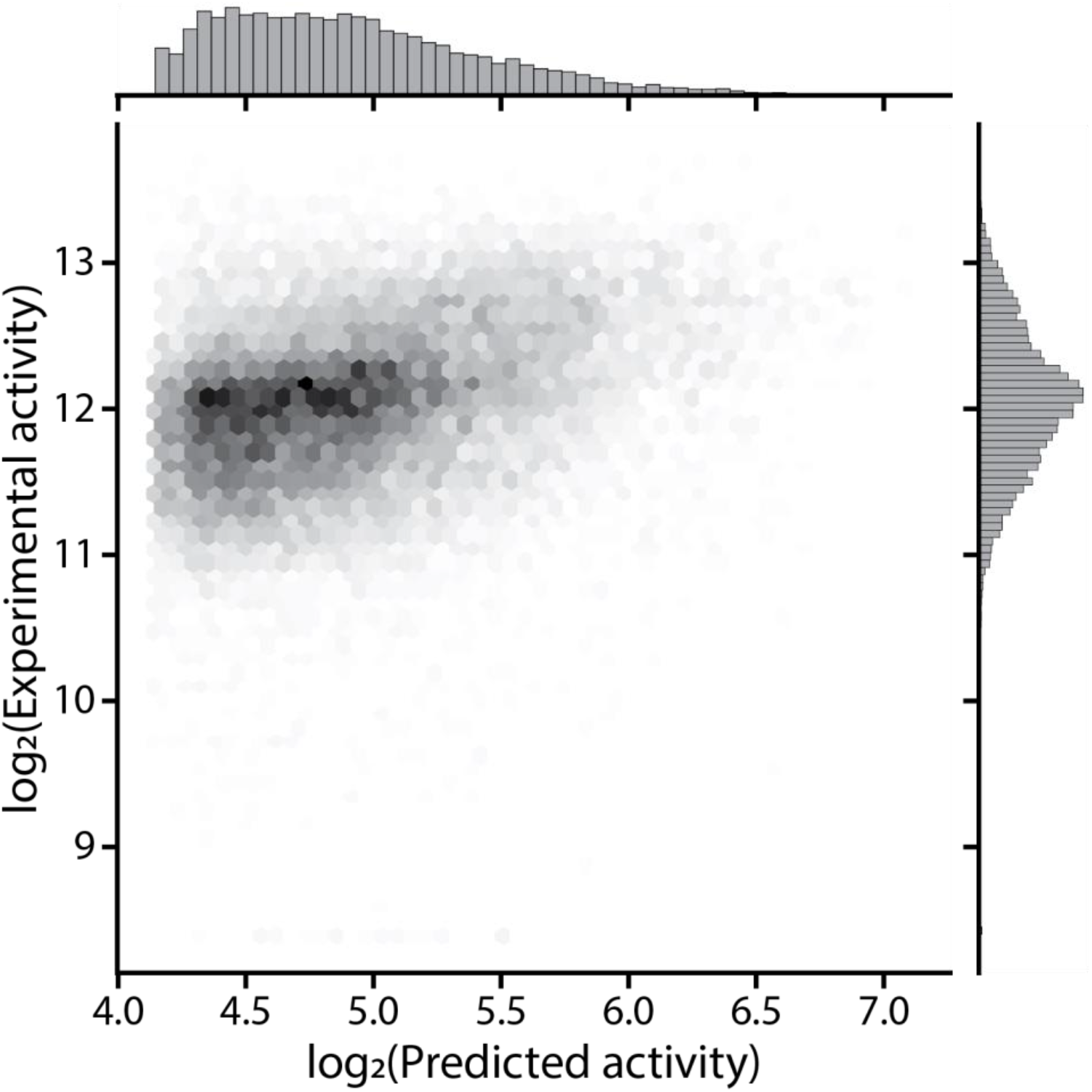
PADDLE predictions are monotonically correlated with experimental observations for combined Arabidopsis and yeast data. Predicted versus measured AD activity of the entire library.

**Figure S5.**
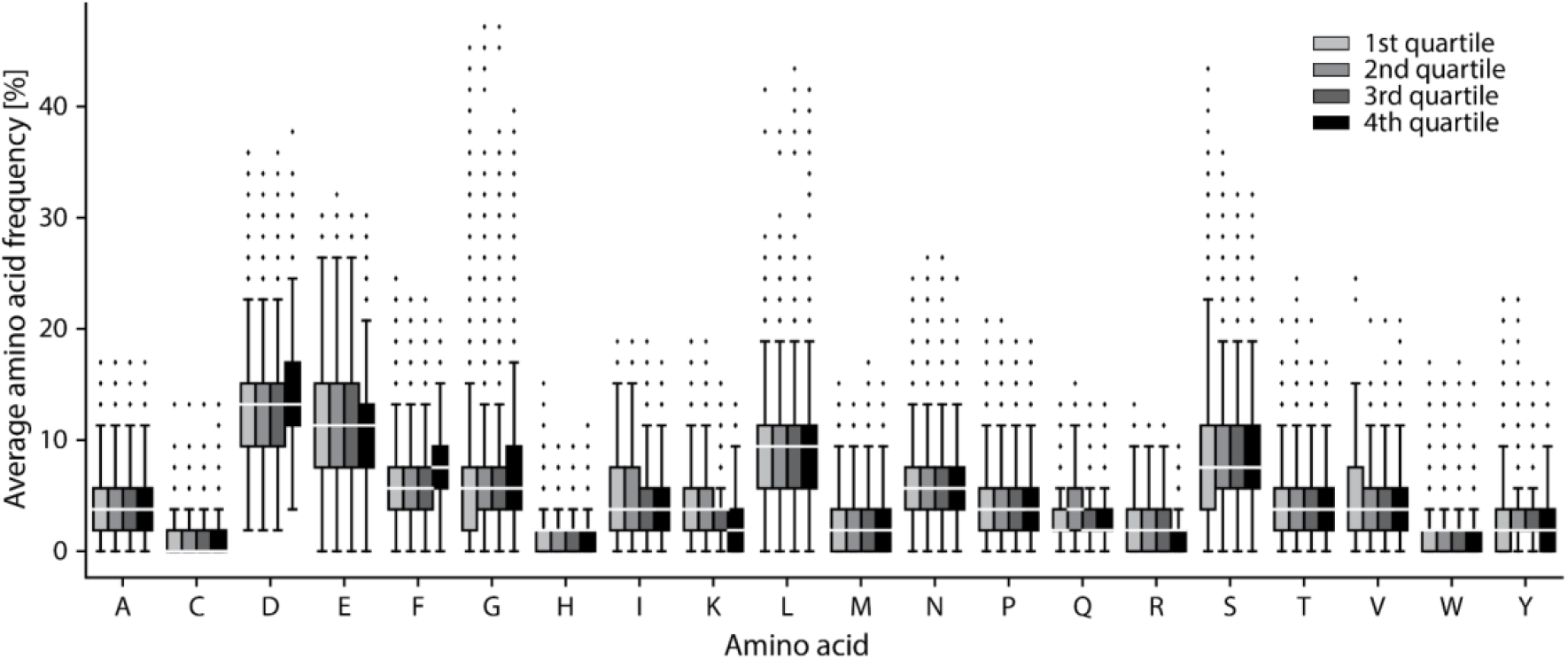
Amino acid composition does not dictate AD activity. Amino acid frequencies of every tile split into quartiles based on activity. Quartile 4 contains the strongest ADs.

**Figure S6.**
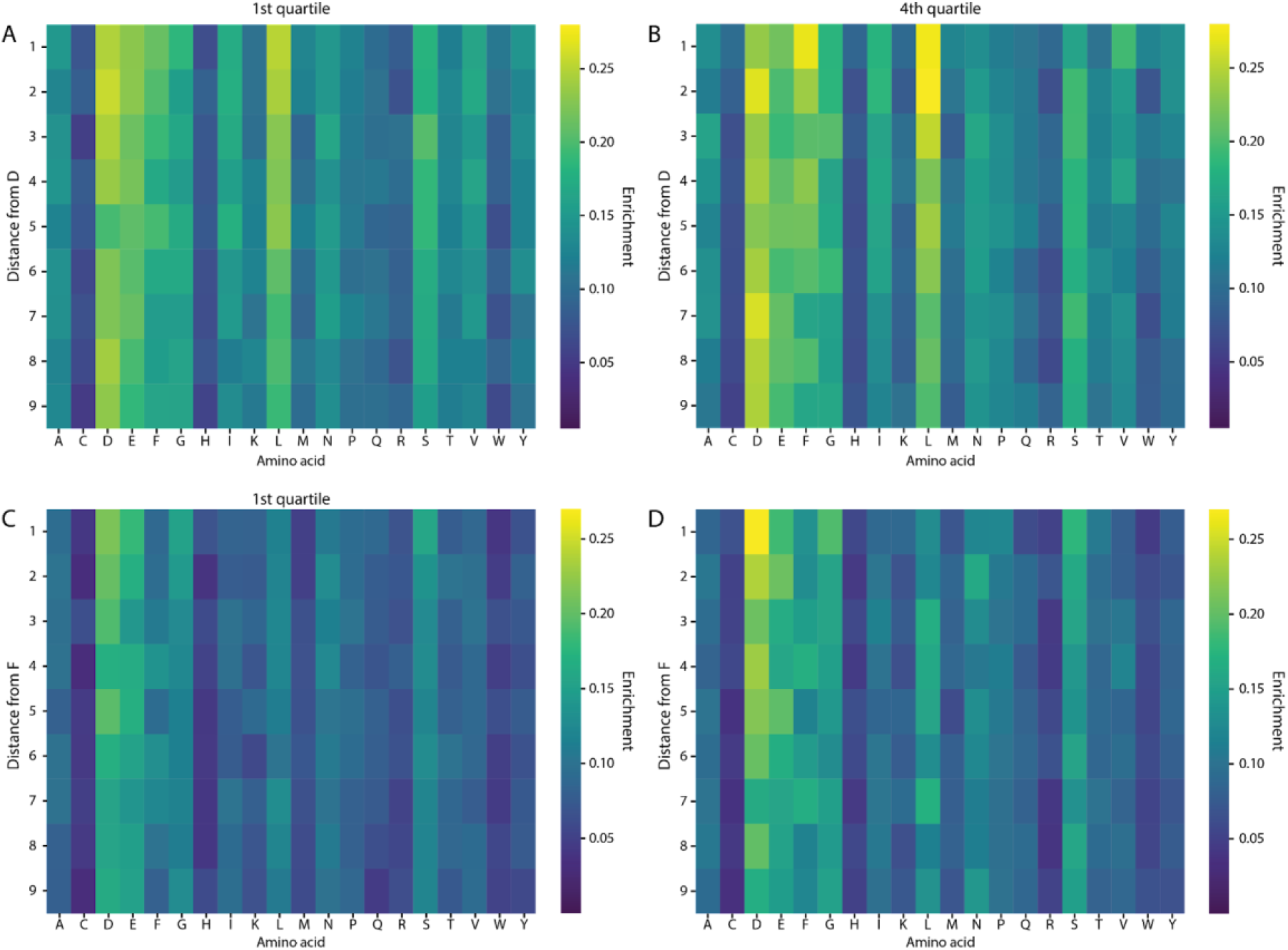
Dipeptide positioning suggests sequence grammar in the strongest quartile. The x-axis in all panels depicts each possible amino acid to pair into a dipeptide. The y-axis describes the distance of the first amino acid to the second amino acid. The heat describes the enrichment of dipeptides. As an example in (**A**), a distance of two on the y-axis to A on the x-axis represents the enrichment of AxD/DxA dipeptides. We performed this analysis using all AAs as the viewpoint, and show the 2 with the strongest signal. D residues frequently have hydrophobic residues F,L,V upstream or downstream when comparing the (**A**) 1st quartile and (**B**) fourth quartile. F residues frequently have acidic residues immediately upstream or downstream when comparing the (**C**) 1st quartile (**D**) 4th quartile.

**Figure S7.**
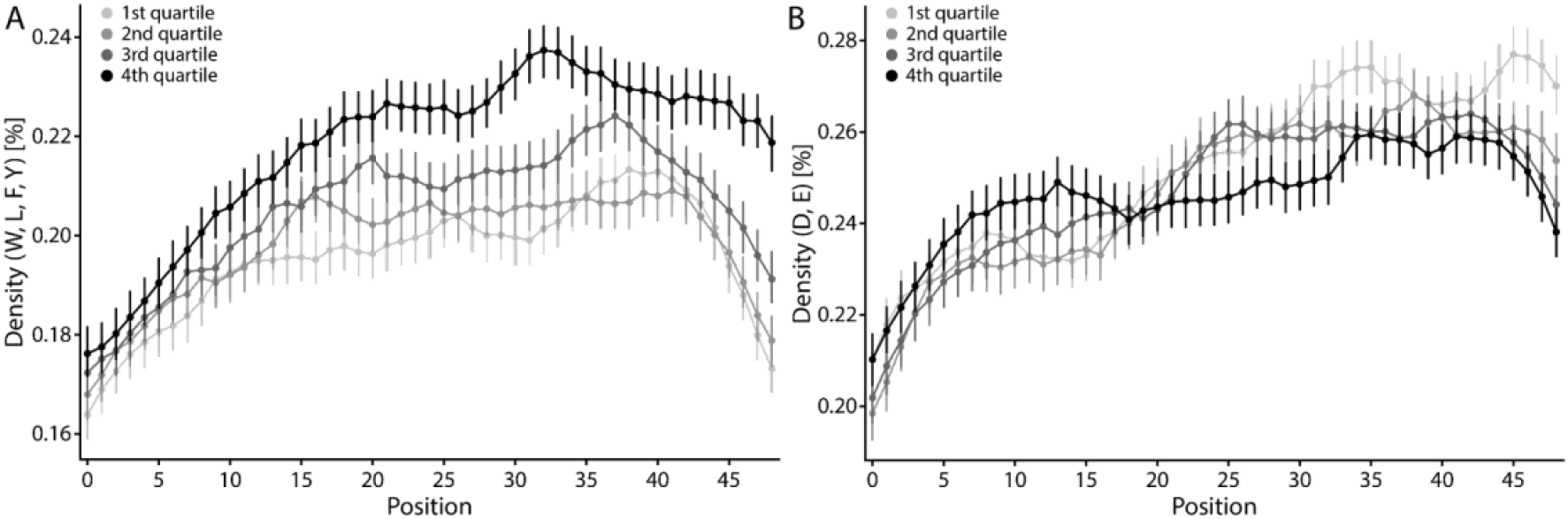
AD populations with similar amino acid composition show disparate amino acid residue distribution. Density of functional amino acids across every position of every tile in the quartile with the strongest and weakest activity (4388 sequences per quartile). Density is calculated in a five amino acid window for each position along the AD as the average of all (**A**) hydrophobic residues (W, L, F, Y), (**B**) acidic residues (D, E), at the respective position. Error bars indicate the 95% confidence interval.

**Figure S8.**
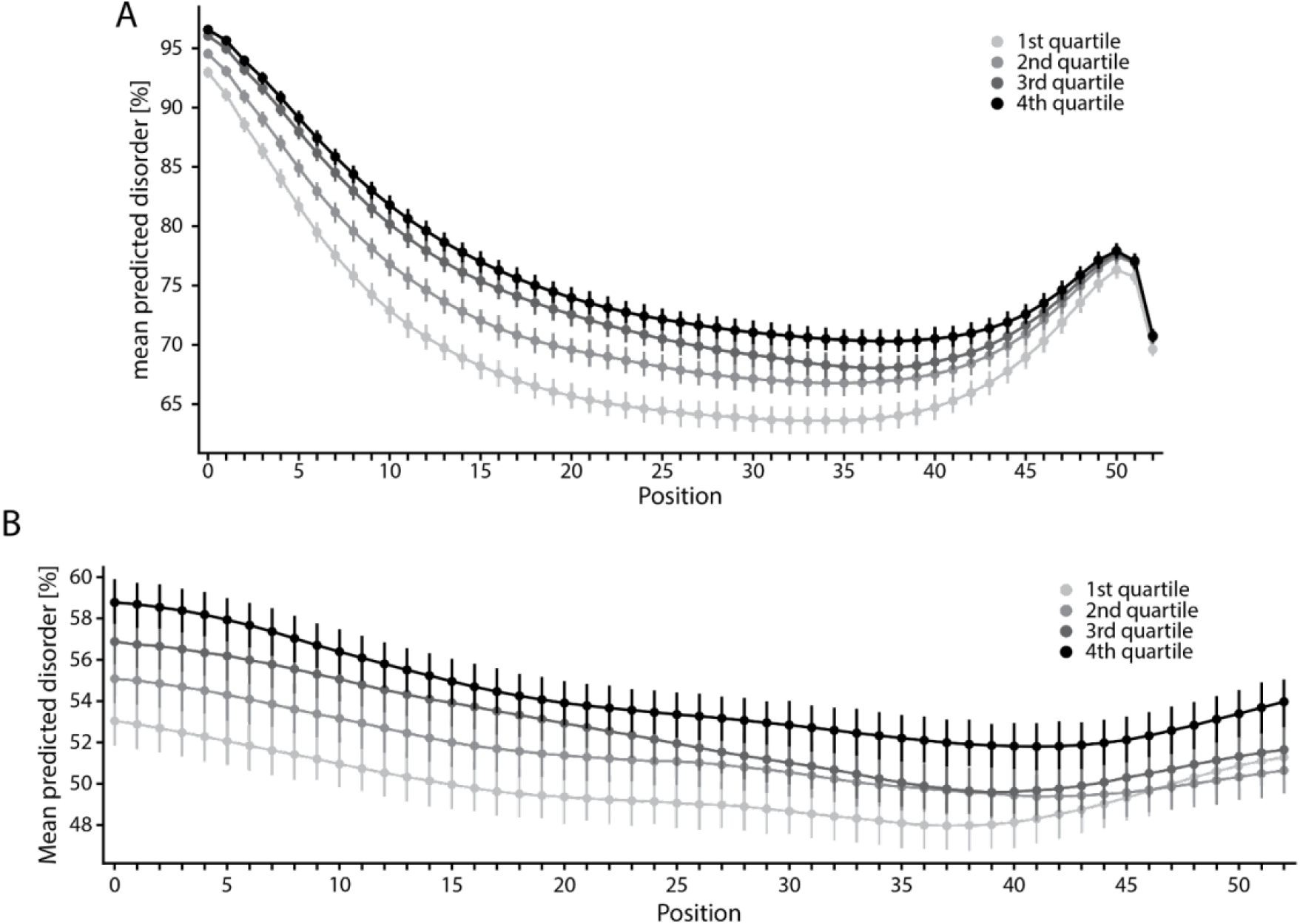
Disorder predictions of tiles in endogenous or synthetic TF context display similar trends. Mean predicted disorder at every position of tiles (**A**) fused to the synthetic TF or (**B**) in their endogenous context in respective quartiles. Error bars indicate the 95% confidence interval. In the synthetic TF, there is increased predicted disorder at the N-terminus from the linker sequence, and increased intrinsic disorder at the C-terminus by virtue of being at the end of the protein.

**Figure S9.**
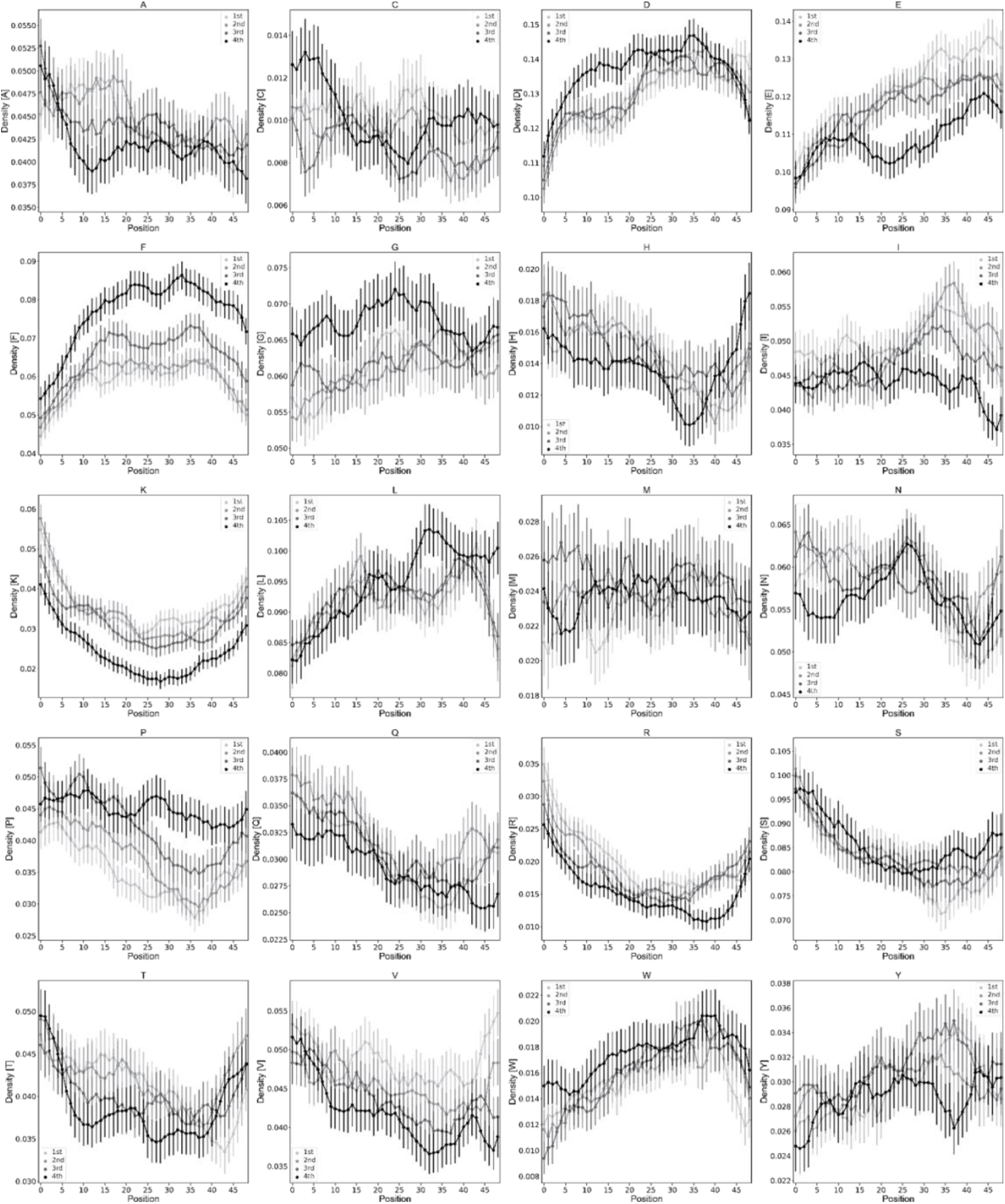
Density of single amino acids varies along tiles between quartiles. Amino acid frequencies in every AD quartile population. Quartile 4 contains the strongest ADs. Density of functional amino acids across every position of every tile in the quartile with the strongest and weakest activity (4388 sequences per quartile). Density is calculated in a five amino acid window for each position along the AD as the average of the respective tile.

**Figure S10.**
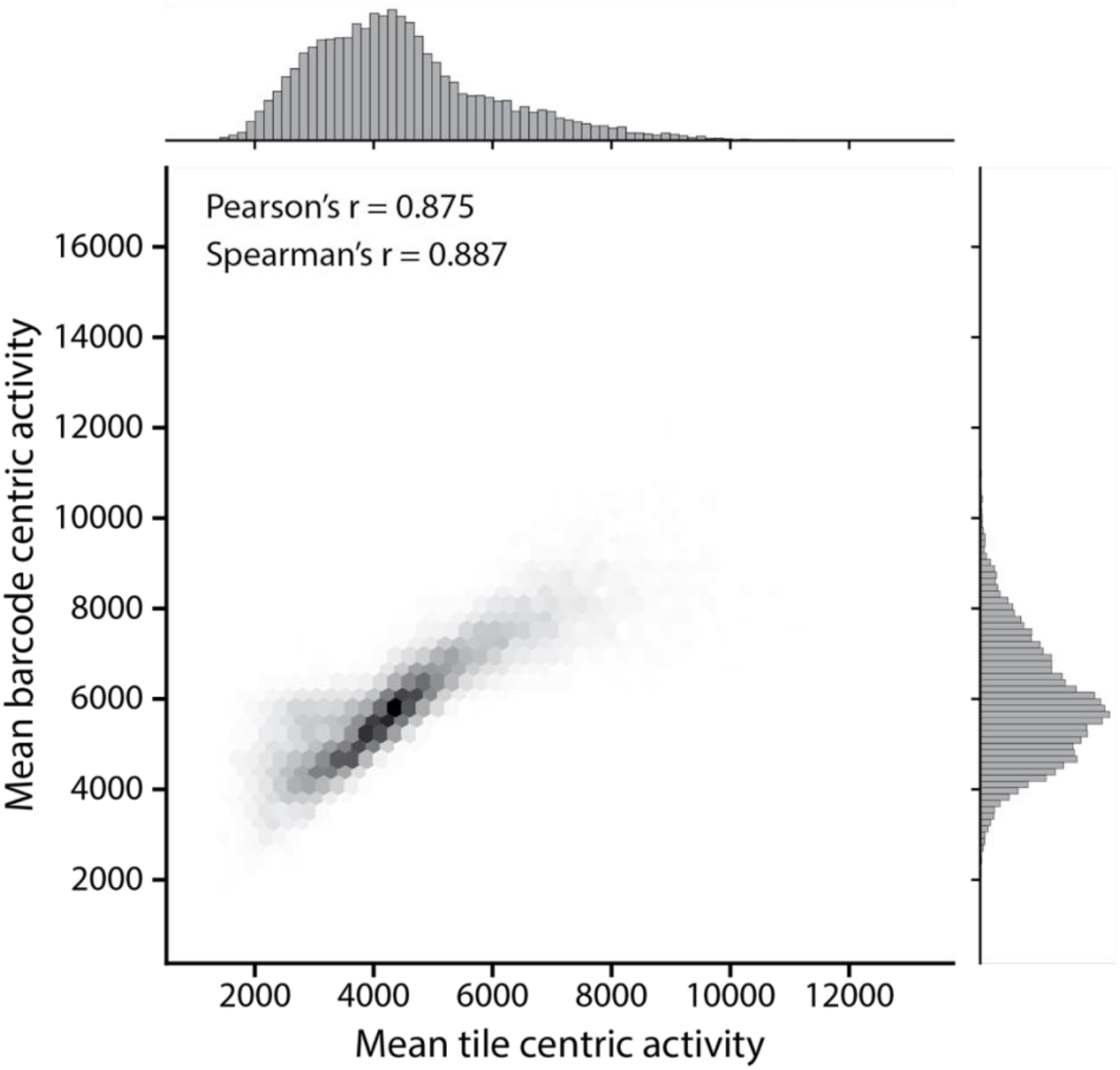
Activity derived from independent barcodes is comparable to activity of pooled barcodes. Mean barcode centric activity was generated by calculating the mean of the activity of independent barcodes of each tile. The mean tile centric activity was derived by pooling the reads in the respective bins of all barcodes associated with a tile and then calculating activity. Shown is the dataset after removing all barcodes less than 1000 reads.

## Notes

### Competing Interest Statement

Pending provisional patent.

